# Propionate oxidation by *Geobacter sulfurreducens* is electron acceptor dependent

**DOI:** 10.64898/2026.02.11.701306

**Authors:** David Hernández-Villamor, Jean-Romain Bautista Angeli, Aya Jeaidi, Andrea Joaquín-García, Korneel Rabaey, Antonin Prévoteau

## Abstract

The accumulation of propionate is a challenge in numerous fermentative industrial processes because its degradation is energetically unfavorable and limited to few microbial species. Here, we report for the first time the oxidation of propionate by the extracellular electron transfer (EET)-capable bacterium *Geobacter sulfurreducens* in axenic cultures. *G. sulfurreducens* was capable of utilizing propionate both as electron donor (ED) and source of carbon with fumarate as electron acceptor (EA). In contrast, propionate was metabolized only in the presence of acetate with soluble Fe(III) citrate, and was not oxidized when insoluble iron oxides or glassy carbon electrodes poised at +0.1 V vs. SHE were the EAs. Biomass yield (per mole of electrons available) was lower with propionate alone than with propionate and acetate together, and acetate was preferentially consumed when both were present. Transcriptomic analysis of cultures grown with either propionate or acetate (with fumarate as EA) showed significant gene expression shifts strongly suggesting the methylmalonyl-CoA pathway as the main route for propionate degradation. Furthermore, propionate-consuming cultures exhibited an upregulation of branched chain amino acids (BCAAs) biosynthesis, as well as sulfur, nitrogen, and 2-oxocarboxylic acids metabolism.

**IMPORTANCE:** The accumulation of propionate is a challenge in anaerobic and fermentative processes because it inhibits methanogenesis, and few microbial species within such systems can degrade it. *G. sulfurreducens* is a model electroactive bacterium widely used in bioelectrochemical systems and is increasingly studied in wastewater treatment and anaerobic digestion because of its ability to enhance syntrophic metabolism via direct interspecies electron transfer. We show for the first time that *G. sulfurreducens* can oxidize propionate, expanding its known metabolic repertoire, and that this capability is controlled by the nature of the terminal electron acceptor. Transcriptomic analyses strongly suggest that the methylmalonyl-CoA pathway is the main pathway for propionate degradation and reveal additional associated transcriptional changes. These findings, together with insights into propionate degradation kinetics, could inform future strategies aimed at using this bacterium to mitigate propionate buildup and improve the stability of anaerobic treatment systems.

## INTRODUCTION

*G. sulfurreducens* is a Gram-negative species well-known for its capability to perform Direct Electron Transfer (DET). In this process, the electrons from substrate oxidative metabolism are transferred throughout the perisplasmic space and the outer membrane to reduce by direct contact solid extracellular electron acceptors (EAs). *G. sulfurreducens* can utilize a wide range of EAs, both soluble (fumarate [1], malate [2], Cr(VI) [3], U (VI) [4], Co (III)-EDTA [2], Fe(III) citrate [2, 5], humic acids [6], anthraquinone-2, 6-disulfonate (AQDS) [6]) and insoluble (Fe(III)- and Mn(IV)-(oxyhydr)oxides [7], elemental sulfur [2], various anode materials [8]). To a lesser extent, it can also respire O_2_ under microaerobic conditions [9, 10]. *G. sulfurreducens* prefers to use acetate as electron donor (ED) [1, 2], yet can also use H_2_ [2, 11], CO [12], formate [1, 13], lactate [1, 14], and cathodes [15]. In addition, acetate and pyruvate are well-known sources of assimilable carbon [11], whereas fumarate can only contribute under specific conditions and less efficiently [2, 11]. While *G. sulfurreducens* is metabolically very versatile, its ability to utilize propionate either as ED or carbon source has never been shown even though it could be of relevance to various applications such as anaerobic digestion (AD) and wastewater treatment, among others.

In AD the accumulation of propionate is considered a central issue that limits its performance [16]. The oxidation of propionate is thermodynamically constrained and only occurs at low H_2_ partial pressures (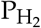 = [1 to 10] Pa) [17, 18]. These values are maintained through tight syntrophic interactions between propionate-oxidizing bacteria and hydrogenotrophic methanogens. The disruption of these cause propionate accumulation, leading to acidification and a decrease of methanogenic activity [19]. The same phenomenon occurs in anaerobic wastewater treatment, where AD-like microbial communities are disrupted, affecting process stability and efficiency [18]. Direct Interspecies Electron Transfer (DIET), a process where different microbial species exchange electrons either through direct contact or via conductive particles, has recently been investigated as an alternative to H_2_-mediated interactions to improve process efficiencies by preventing propionate accumulation [20, 21]. As a DIET-capable bacteria, *G. sulfurreducens* has been co-cultured with *Escherichia coli* for wastewater treatment [22], and with *Syntrophobacter fumaroxidans* and *Syntrophomonas wolfei* in a microbial fuel cell system for propionate and butyrate oxidation to be used in AD, respectively [23, 24]. Furthermore, the introduction of a natural or synthetic microbial consortia to achieve organic acids removal, including propionate, and complete CO_2_ recovery from organic waste is being actively investigated in regenerative life support systems using microbial electrochemical technologies [25]. For these, the introduction of *G. sulfurreducens* is of special interest considering its metabolic versatility, capacity to form electroactive biofilms, and broad use in bioelectrochemical systems [26].

In contrast to *G. sulfurreducens*, both *G. metallireducens* and *G. anodireducens* are known to degrade propionate. In *G. metallireducens*, this likely occurs via the methylcitrate or methylmalonyl-CoA (MMC) pathways [27, 28], whereas in *G. anodireducens*, propionate oxidation is associated with the MMC pathway [29].

In this work, we investigated the capabilities of axenic *G. sulfurreducens* cultures to oxidize propionate when supplied as single ED or in combination with acetate, under four different EAs, namely fumarate, soluble and insoluble forms of Fe(III) and glassy carbon electrodes poised at +0.1 V vs. SHE, a potential at which the metabolic energy yield per electron delivered is maximized, thus minimizing any thermodynamic constraints [30]. After showing that *G. sulfurreducens* can use propionate with specific EAs, the gene expression profile of acetate and propionate grown cultures using fumarate as EA was quantified via RNAseq, then compared to elucidate the putative metabolic pathways active during propionate oxidation. Both differential gene expression (DGE) and biological enrichment analyses were performed to obtain further metabolic insights linked to the consumption of propionate in contrast to acetate.

## MATERIALS AND METHODS

### *G. sulfurreducens* pre-growth

*G. sulfurreducens* PCA (DSMZ 12127) was cultured at 28°C and 100 rpm agitation speed in penicillin flasks using modified M9 medium containing 6 g L^−1^ Na_2_HPO_4_ · 2 H_2_O, 3 g L^−1^ KH_2_PO_4_, 0.5 g L^−1^ NaCl, 0.5 g L^−1^ NH_4_Cl, 0.1 g L^−1^ MgSO_4_ · 7 H_2_O, and 0.014 g L^−1^ CaCl_2_ · 2 H_2_O (all reagents were supplied by ChemLab and Carl Roth unless otherwise specified). The solution was then sparged with N_2_ (Air Liquide) for 25 min and 0.5 g L^−1^ of NaHCO_3_ was added before sealing the flasks with rubber stoppers and metal clamps. The head space was flushed with a mix of N_2_:CO_2_ gas (90:10 v/v, Air Liquide) for 15 min and the pressure kept at 130 kPa before autoclavation. The pH was adjusted to 7.4 using 1M NaOH, and the media was supplemented with a mix of trace elements and vitamins [31]. Acetate (10 mM) was added as carbon source and ED and fumarate (40 mM) as EA. To avoid a carry-over of residual volatile fatty acids (VFAs), full depletion of either VFA, or both, was ensured for each experiment on a case-by-case basis using ion chromatography (IC) (see Supp. Materials and methods). Cells at the beginning of stationary phase were used as inoculum for subsequent experiments.

### Growth assays

The use of propionate as ED, either alone or in combination with acetate, was assessed in penicillin flasks (150 mL total volume) using three EAs respectively, namely fumarate, soluble Fe(III) citrate and insoluble Fe(III) oxides. Flasks were inoculated with either *G. sulfurreducens* or 0.22 µm-filtered spent medium (5% v/v), to serve as sterile control (unless otherwise specified).

A summary of the replicates utilized per experimental condition is provided in Table S1. Cultures were grown at 28°C in the dark and 150 rpm agitation speed. *G. sulfurreducens* cultures were confirmed axenic via Sanger sequencing (see Supp. Materials and methods). Flasks were sampled regularly to monitor VFAs, Fe(II) and total dissolved iron.

Standard redox potentials and electron stoichiometry for propionate and acetate oxidation with either fumarate or Fe(III) citrate utilized to obtain changes in free energy estimates are provided in Table S2.

A ferrozine assay (ref. 91836, Macherey-Nagel) was performed to quantify Fe(II) and total iron [32]. Extracted samples were immediately diluted in HCl (0.15 M) to prevent Fe(II) oxidation and subsequent precipitation. Cell density was quantified via flow cytometry (see Supp. Materials and methods) at the beginning and end of the experiments containing soluble EAs.

#### Fumarate reduction assays

Flasks contained M9 medium with fumarate (60 mM) as EA, and either propionate (4 mM) as only putative ED, or both propionate and acetate (4 mM each).

#### Fe(III) citrate reduction assays

Ammonium Fe(III) citrate (60 mM) (18% Fe w/w, Sigma-Aldrich) was used as EA in modified M9 medium. However, the buffer was modified to 2 g L^−1^ NaHCO_3_ and 0.069 g L^−1^ NaH_2_PO_4_ · H_2_O ([33]) to minimize iron-phosphate precipitation (see Fig. S1). One experimental condition contained only propionate (2 mM) and the other contained propionate and acetate (2 mM each).

#### Solid Fe(III) oxide reduction assays

A 25% v/v inoculum was added in modified M9 medium containing the less concentrated phosphate buffer. The Fe(III) oxides were prepared dissolving 6 g of FeCl_2_ in 25 mL deionized water (DI), adjusted to pH 8.0 using 1 M NaOH. Next, 30% w/w H_2_O_2_ was added drop-wise until the solution changed color from green to reddish-brown, which indicates the formation of Fe(III) oxide precipitate. This was collected by centrifugation (7,100 g) for 20 min and washed with DI water, then dried in an oven at 105°C overnight. All flasks, containing the less concentrated phosphate buffer, were supplemented with Fe(III) precipitates accounting for 45 mM of Fe(III) equivalents. Conditions tested included propionate (2 mM) alone or propionate with acetate ([2 and 4] mM, respectively).

### Electrochemical experiment

Laboratory bottles (GL45, 100 mL working volume, Glasgeratebau Ochs, Germany) served as single-compartment electrochemical cells. Each bottle, containing a magnetic stirring bar, was sealed using a GL45 polybutylene terephthalate open cap and rubber stopper containing a sampling port, a graphite rod (Morganite Luxembourg) as counter electrode, a glass reference electrode holder, and a planar glassy carbon working electrode (1 cm^2^, HTW, Germany), previously polished using alumina slurries (ALS, Japan) of decreasing particle sizes ([3, 1 and 0.05] µm) on microcloth pads and rinsed with DI water. The fully assembled electrochemical cell, devoid of medium, was sterilized by autoclaving and subsequently transferred to a biosafety cabinet. Then, anaerobic modified M9 medium (pH 7.4) was added by filter sterilization, and a Ag/AgCl 3 M KCl reference electrode (ALS, Japan, + 0.205 V vs. SHE at 28 °C) was assembled into the holder, which also contained 3 M KCl. The electrochemical cells were exposed to UV-light for 20 min and flushed with N_2_:CO_2_ (90:10 v/v) through a 0.22 µm-filter to maintain sterility and ensure anaerobic conditions. Next, the cells were introduced into an anaerobic chamber (GP-Campus, Jacomex, TCPS NV, Rotselaar, Belgium) maintained at 28 °C and fed with a gas mixture of N_2_:CO_2_ (90:10 v/v).

The electrochemical cells were set on a stirring plate (Poly15, Thermo Fisher Scientific) at 130 rpm (setup in Fig. S2). The rubber stoppers were sprayed with ethanol 70% v/v before inoculating *G. sulfurreducens* (8% v/v) pre-grown in the same medium using acetate as ED and fumarate as EA, both depleted before inoculation. Acetate (7 mM) was added to one system to serve as a control, and propionate (4 mM) was tested in triplicate. Electrochemical measurements were obtained using a VSP potentiostat (Biologic, France). All potentials are referenced to standard hydrogen electrode (SHE). The working electrode, serving as EA, was set to +0.1 V vs. SHE, which lies on the anodic plateau of *G. sulfurreducens* catalytic cyclic voltammetry profiles [31], thus maximizing metabolic energy gains per electron delivered to the electrode.

### RNA sequencing and analysis

#### Sample preparation

*G. sulfurreducens* was cultured with fumarate (60 mM) as EA and either acetate (7 mM) or propionate (4 mM) as sole ED and carbon source (n = 4 replicates per condition). Biomass samples (4 mL each) were collected when approximately 75% of the supplied electron donor had been consumed and their consumption rate was maximal. Additionally, in the acetate-only condition, propionate was added upon complete depletion of acetate to induce a metabolic transition to propionate utilization (“shift” condition, n = 4). Cell pellets were obtained by centrifugation at 4°C and 19,000 g for 5 min, then resuspended in 300 µL RNAlater^®^ (Sigma-Aldrich) and incubated overnight at 4°C. Next, the samples were centrifuged again with the same settings, and the supernatant was discarded before the pellets were stored at –80°C.

#### RNA extraction, sequencing and alignment

RNA extraction, quality control, library preparation, sequencing, and genome alignment were outsourced to BaseClear B.V. (Leiden, Netherlands). Total RNA was extracted from 12 bacterial cell pellets, and RNA integrity and potential DNA contamination were assessed prior to library preparation. An input of 200 ng of RNA was used, and rRNA was depleted using the Illumina Ribo-Zero Plus rRNA depletion kit. The mRNA libraries were prepared using the Illumina protocol and quality-checked before sequencing. Libraries were sequenced on an Illumina NovaSeq 6000 platform to generate 150 bp paired-end reads, with an average yield of 3 GB per sample.

Raw reads were demultiplexed and converted to FASTQ format using bcl-convert v4.2.4 (Illumina). Low-quality bases and adapter sequences were removed using fastp v0.23.4 [34], and quality control was assessed with FastQC v0.11.9 [35]. Reads containing PhiX control sequences were filtered out using Bowtie2 v2.4.5 [36].

For alignment, quality-trimmed reads were mapped against the reference genome (GCA_000007985.2) using STAR v2.7.1 [37]. Mapped reads were assigned to genomic features using featureCounts v1.6.3 [38]. Gene expression counts were normalized using the variance stabilizing transformation (VST) method from the R package DESeq2 (v 1.44.0) [39].

#### Differential gene expression and enrichment analyses

DGE between the propionate and acetate conditions was performed using DESeq2 on raw count data [39]. Genes with an absolute log_2_ fold change > 1 and an adjusted p-value < 0.05 were considered significantly differentially expressed.

These gene sets were subsequently used for over-representation analysis (ORA) using the KEGG database with the clusterProfile (v 4.12.6) R package [40]. Since the KEGG annotation for *G. sulfurreducens* in the official database was outdated, KEGG orthologs were reassigned using EggNOG-mapper (v2), which provides functional annotations based on orthology with updated reference data from the EggNOG database [41].

#### Propionate signature score

Gene set variation analysis (GSVA) was performed using the GSVA R package (v 1.52.3) to calculate a propionate signature score [42]. This gene signature was defined by 10 candidate genes: GSU1103, *ackA, pta, ato-1, ato-2*, GSU3299, GSU3300, *mceE*, GSU1578, and GSU3302. Variant stabilized transformation (VST) normalized expression data was used as input for the analysis.

#### Statistical analysis

Comparisons between the three experimental conditions were performed using the Kruskal–Wallis test. False discovery rates (FDRs) were calculated using the Benjamini–Hochberg method to correct for multiple testing, where applicable [43]. This correction was applied to both the differential expression results (DESeq2) and the KEGG pathway enrichment analysis. The P-values and FDR-adjusted p-values below 0.05 were considered statistically significant. All statistical analyses were conducted in R (version 4.4.1).

## RESULTS AND DISCUSSION

### *G. sulfurreducens* consumes propionate with fumarate as EA

The ability of *G. sulfurreducens* to consume propionate as its sole ED and source of carbon (Fig. 1A and B) was evaluated using fumarate as EA. During the first 24 h, the concentration of propionate decreased slightly, from (3.8 ± 0.1 to 3.3 ± 0.0) mM. In the following hours, the consumption rate accelerated so that 75% of the initial concentration was consumed by *t* = 69 h, and full depletion was confirmed at *t* = 93 h (see experimental flasks in Fig. S3). During this period, fumarate decreased by 20.5 mM from its initial value of 52 mM. The major fraction of this decrease (15.2 mM) was microbially reduced to succinate and ∼5 mM was hydrated to yield malate, following Eqs. 1 and 2, respectively. After propionate was depleted, the concentration of succinate plateaued, while fumarate was further converted to malate, reaching (28.8 ± 0.7) mM at 214 h.

**FIG 1.**
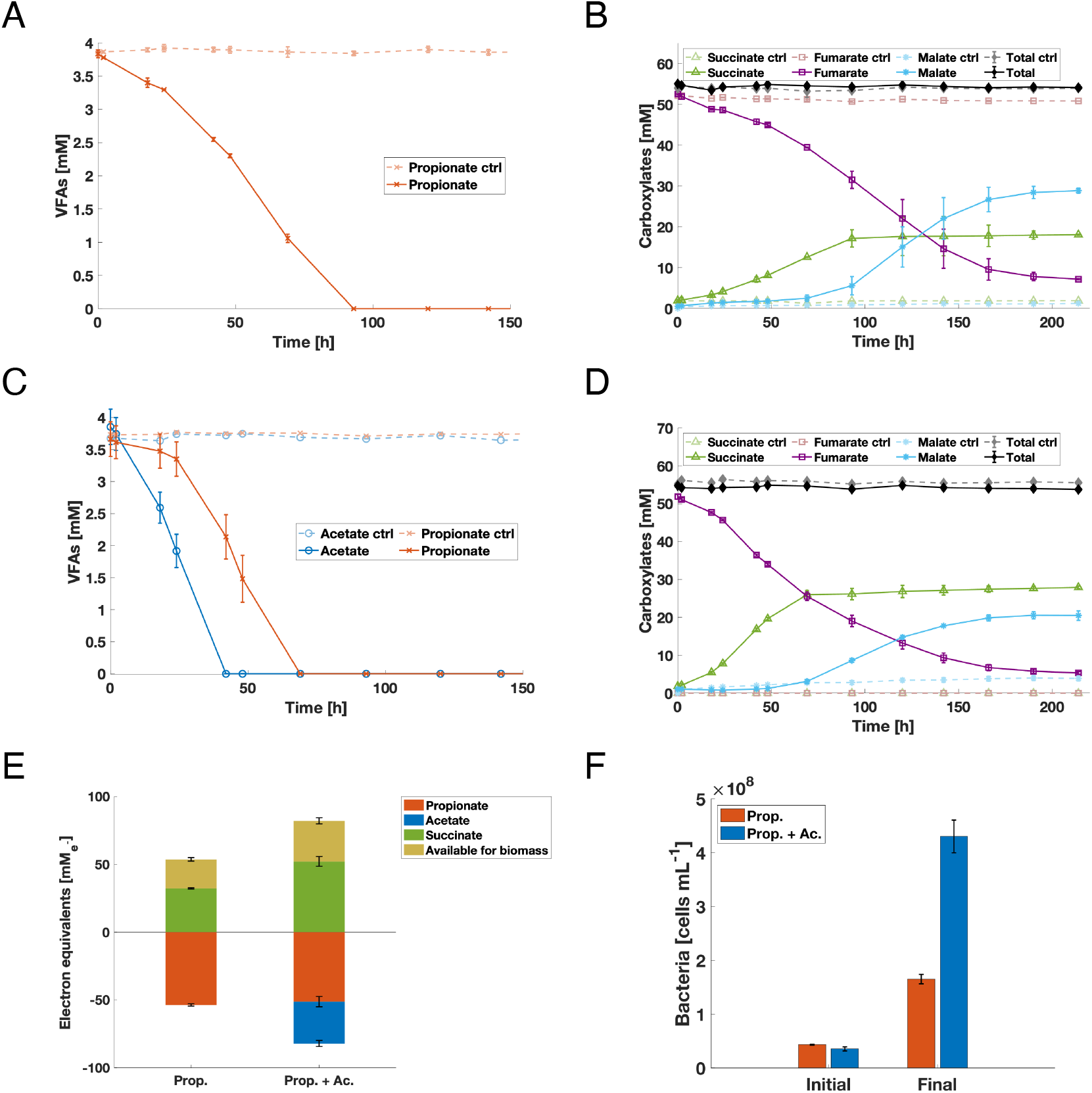
Propionate consumption assay with *G. sulfurreducens* using fumarate as EA. Sterile controls and biological experiments are represented with discontinued and solid lines, respectively. **(A** and **B)** Only propionate is provided as the carbon source and ED. The first shows the evolution of propionate concentration, and the second, the concentration of fumarate and carboxylates formed from its conversion. **(C** and **D)** Same as above with both propionate and acetate initially supplied. **(E)** Electron balance at the end of the experiments: negative values are the electrons theoretically available from the oxidation of EDs to CO_2_ (8 per acetate and 14 per propionate; positive values are the electrons dedicated to EA reduction (2 per fumarate). The difference is assumed to represent the fraction of electrons directed to biomass growth. **(F)** Cell densities at the beginning and end of the experiments.

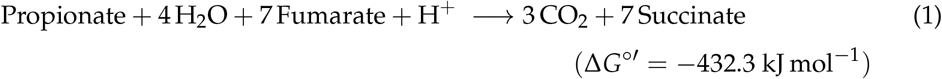

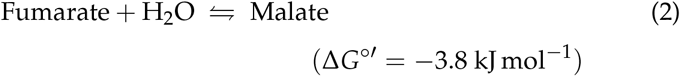

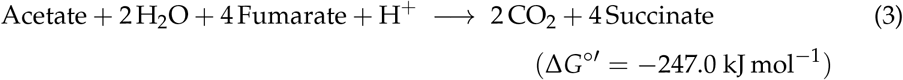

The competition between acetate and propionate as ED was also examined (Fig. 1C and D). Starting at (3.9 ± 0.3) mM for both VFAs, acetate decreased by 50% in the first 24 h following Eq. 3, while propionate decreased by only 10%, confirming that acetate is the preferred ED for *G. sulfurreducens*. Upon acetate depletion (*t* ≈ 42 h) the rate of propionate consumption increased until full depletion by *t* ≈ 69 h. Similarly to the propionate-only condition, fumarate was first reduced to succinate until all EDs were consumed, then it was slowly converted to malate. The capacity of *G. sulfurreducens* to utilize fumarate as carbon source by promoting gluconeogenesis has been described under ED-limiting conditions [11], yet the sum of the concentration of fumarate, succinate and malate remained constant in our work (Fig. 1B and D).

An electron balance was performed for each experimental condition (Fig. 1E) assuming complete EDs oxidation into CO_2_. Electrons supplied are reported as negative values while electrons recovered in succinate are positive. The electrons not recovered in succinate (40% in the propionate cultures, and 36.5% in the propionate plus acetate cultures) were likely diverted to biomass growth (cell synthesis and exopolysaccharides formation). These values are in line with previous studies, which report a 27.5% to 45% of equivalent electrons toward cell synthesis [44, 45, 11]. The evolution of the cumulative fraction of electrons assumed to go to biomass was monitored, exhibiting a similar trend in both conditions, yet with propionate alone being slightly higher (Fig. S4A). Furthermore, the fraction of electron equivalents not recovered in measured reduced end-products was evaluated for each time interval between two sampling points (Fig. S4B), showing stable yield at ∼40% when the culture first oxidized predominantly acetate then only propionate.

Cell concentrations were assessed in both experimental conditions (Fig. 1F). When propionate and acetate were supplied together, (4.0 ± 0.2) × 10^8^ cells mL^−1^ were produced. This is a 3.1-fold greater increase than with propionate alone, which produced (1.3 ± 0.1) × 10^8^ cells mL^−1^, and is partly explained by the concentration of initially available electron equivalents. This increase in total biomass caused propionate to be depleted ∼24 h earlier in the propionate plus acetate condition with respect to propionate supplied alone.

To estimate a biomass synthesis yield, the concentration of cells was normalized to the moles of electrons available to produce biomass. The propionate plus acetate condition yielded (1.3 ± 0.1) × 10^10^ cells 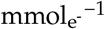. This is a 2.3-fold increase in biomass production with respect to the propionate-only condition, which yielded (5.7 ± 0.8) × 10^9^ cells 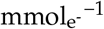. This suggests that *G. sulfurreducens* conserves more efficiently energy from acetate, allowing to direct more energy toward biomass synthesis. These results show, for the first time, the utilization of propionate by *G. sulfurreducens* not just as ED but also as a carbon source when fumarate is present as EA.

### *G. sulfurreducens* consumes propionate with Fe(III) citrate as EA only when acetate is supplied

The consumption of propionate by *G. sulfurreducens* was investigated using soluble Fe(III) citrate both in the absence (Fig. 2A and B) and presence of acetate (Fig. 2C and D). Propionate was not consumed when supplied as only VFA. By contrast, when combined with acetate (each at ∼ 2 mM), acetate was preferentially oxidized and fully depleted within 24 h, and the concentration of propionate declined by 90% at *t* = 48 h. More acetate (∼ 2 mM) was added to try to stimulate further propionate consumption, leaving a small fraction unconsumed after 72 h (see experimental flasks in Fig. S5). In total, 3.3 mM of acetate and 1.85 mM of propionate were oxidized, equivalent to 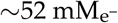 according to Eqs. 4 and 5. This concentration is ∼25% higher than that of recovered Fe(II) ([39.3 ± 1.7] mM), a difference likely representing the fraction of VFAs used for biomass synthesis.

**FIG 2.**
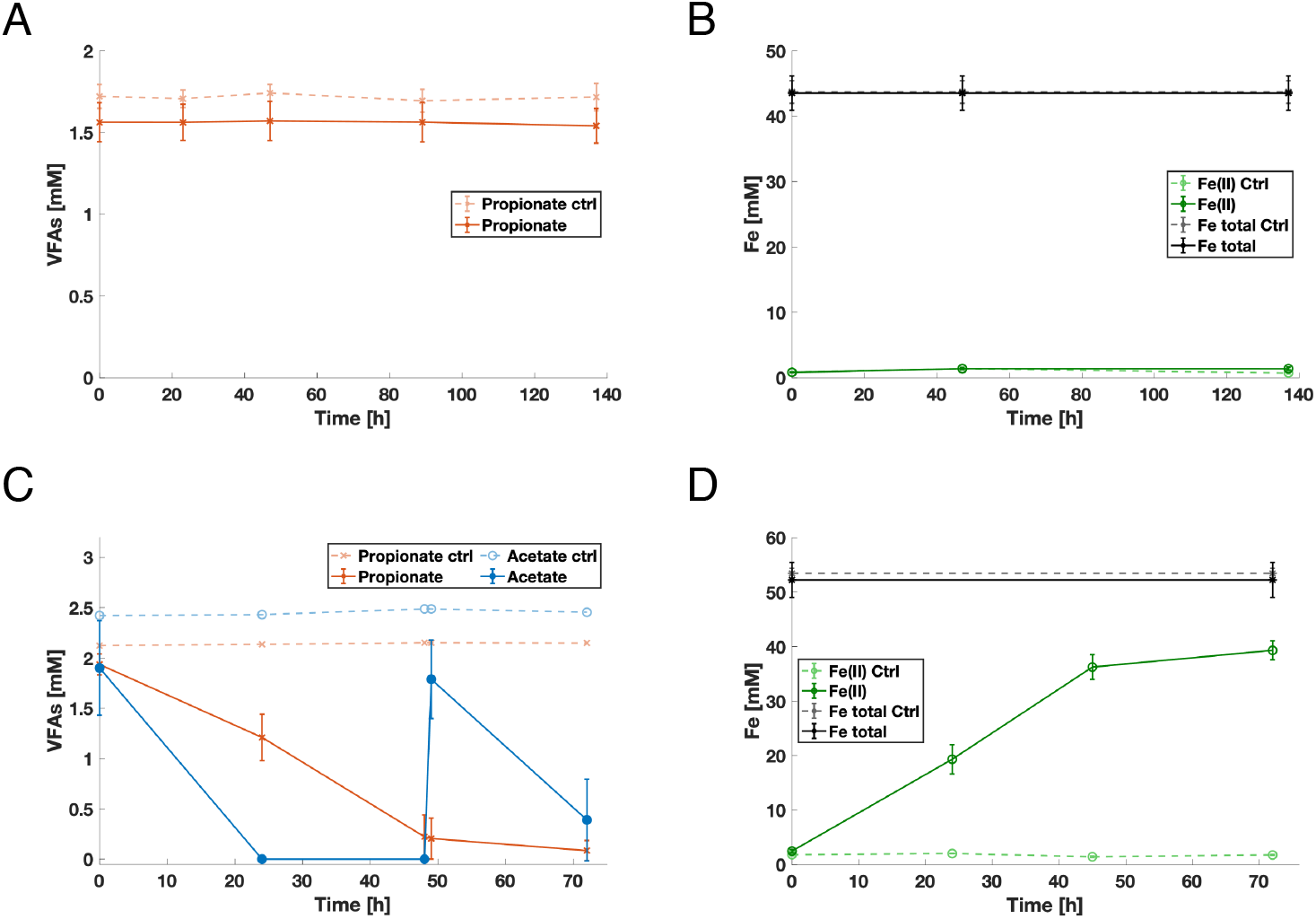
Propionate consumption assay in *G. sulfurreducens* using soluble Fe(III) citrate as EA. **(A** and **B)** Only propionate is provided as the putative carbon source and ED. The first shows the evolution of propionate concentration while the second exhibits the concentration of Fe(II) and total dissolved iron. **(C** and **D)** Same as above. Both propionate and acetate are initially supplied with a second addition of acetate (2 mM) at *t* = 48 h.

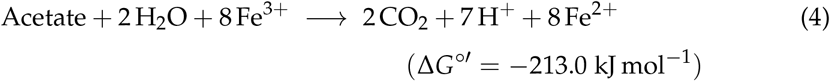

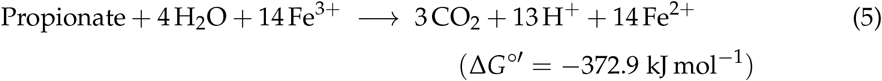

In the acetate plus propionate cultures with soluble Fe(III) citrate, the concentration of cells increased from (0.3 ± 0.1 to 2.4 ± 0.6) × 10^8^ cells mL^−1^ by the end of the experiment (see Fig. S6). In comparison, the corresponding fumarate-respiring cultures supplied with both VFAs produced 4.0 × 10^8^ cells mL^−1^. When normalized by supplied electron equivalents, the biomass yield was 4.0 × 10^9^ cells 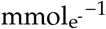 with Fe(III) citrate and 9.1 × 10^9^ cells 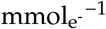 with fumarate. As such, the Fe(III) citrate condition achieved ∼45% of the biomass yield generated with fumarate. Besides, the initial acetate:propionate molar ratio between both experiments is not consistent (1:1 vs. 2:1 with fumarate and iron citrate, respectively), likely underestimating the yield difference observed. Consistently, biomass yields are reportedly 2-to 3-fold lower with Fe(III) citrate relative to fumarate as the EAs, when acetate is used as ED [46, 11].

These results are consistent with previous reports showing that *G. sulfurreducens* conserves less energy when reducing extracellular EAs, even though complexed Fe(III) citrate has been reported with a similar formal potential (*E*^′^, further see Fig. S7). Accordingly, thermodynamic estimates based on standard (Δ*G*^°′^) and real initial experimental conditions 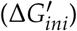 were derived from Eq. 6 (Table S3), indicating that propionate oxidation coupled to fumarate reduction yields more energy per mol of electrons transferred 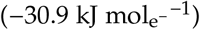 than when coupled to Fe(III) citrate reduction 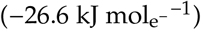, considering Δ*G*^°′^ values.

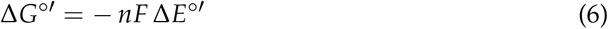

where *n* is the number of electrons transferred, *F* is the Faraday constant (here, 96.485 kJ mol^−1^V^−1^), and Δ*E*^°′^ (V) is the difference between the standard transformed redox potentials of the EA and ED.

Supporting these values, both modeling and experimental studies have shown that *G. sulfurreducens* conserves less energy during Fe(III) citrate respiration (∼0.5 ATP mol_acetate_^−1^), with respect to fumarate ∼1.5 ATP mol_acetate_^−1^ [47, 12].

The requirement for acetate to enable propionate oxidation under Fe(III) citrate–reducing conditions is intriguing. Compared to acetate, propionate degradation is typically less energetically favorable due to several intermediate reactions requiring an energy input [18, 28]. In the comparatively lower energy context of Fe(III) citrate respiration versus fumarate, acetate can potentially provide sufficient energy to prime the propionate oxidation pathway, shift the redox balance, or provide pools of the tricarboxylic acid cycle (TCA) intermediates that are shared between acetate and propionate degradation routes.

### *G. sulfurreducens* does not consume propionate when solid Fe(III) oxides are the EA

The ability of *G. sulfurreducens* to oxidize propionate with precipitated insoluble iron oxides as EA was evaluated with propionate as sole ED (Fig. 3A), or with propionate and acetate (Fig. 3B). No consumption of propionate was detected in either condition. In contrast, acetate was slowly consumed over a total of 76 days, from (3.9 ± 0.6 to 0.2 ± 0.2) mM (see experimental flasks in Fig. S8).

**FIG 3.**
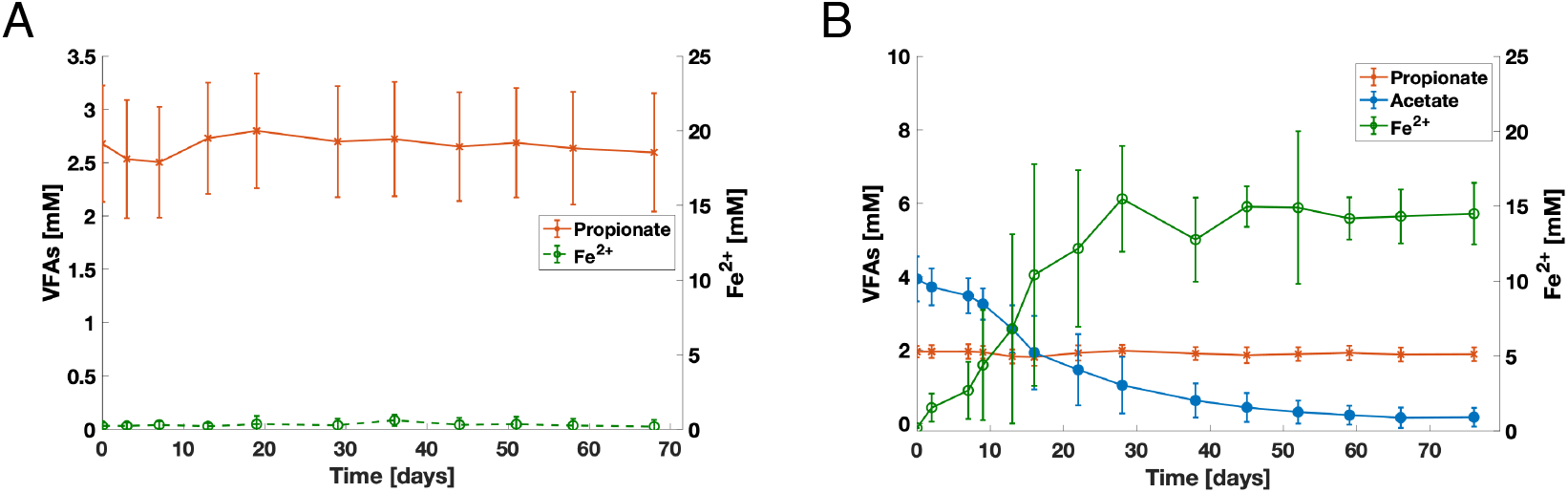
Propionate consumption assay in *G. sulfurreducens* using insoluble Fe(III) oxide as EA. **A)** Propionate as the sole substrate. **B)** Propionate and acetate supplied together. In both conditions, the concentration of VFAs and reduced iron were monitored over time.

The concentration of total Fe(II) formed increased quasi-proportionality to acetate consumed. Yet, only (14.5 ± 2.1) mM of Fe(II) was measured at the end of the experiment, accounting for 45% of the 32 mM of electron equivalent supplied by the acetate. This suggests that the biggest fraction of electrons available was diverted to biomass upon the early biofilm formation on solid EA, as previously reported for *G. sulfurreducens* biofilms grown on electrodes [48]. This time, no accurate cell count measurements could be obtained due to biofilm formation on the iron oxide particles surface.

### *G. sulfurreducens* could not degrade propionate with electrodes as EA

*G. sulfurreducens* was cultured in three-electrode electrochemical cells supplemented with either acetate as positive control (n = 1), or propionate (n = 3) (Fig. 4). Glassy carbon working electrodes were poised at +0.1 V vs. SHE, a potential at which the thermodynamic EA limitation is minimal since all charge carriers are oxidized at the biofilm-electrode interface [31].

**FIG 4.**
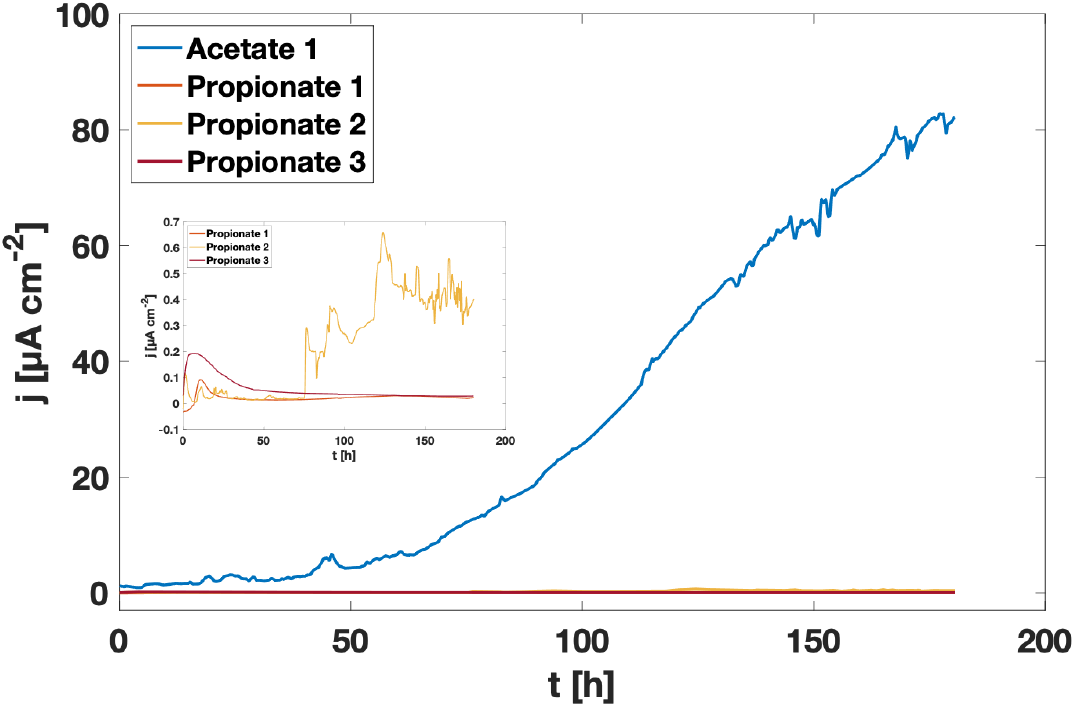
Chronoamperograms from *G. sulfurreducens* cultures with acetate or propionate as ED and glassy carbon electrodes (poised at +0.1 V vs. SHE) serving as EA. Inset: zooms over the propionate-fed replicates.

With acetate, current density first increased slowly during a lag phase characterized by electrode colonization. Thereafter, current increased faster until 82.7 µA cm^−2^ at 178 h. Both the evolution and the magnitude of the current density recorded are consistent with previous works in axenic *G. sulfurreducens* grown with acetate [49, 50, 31].

With only propionate as putative ED, current only slightly increased for two replicates during the first 13 h, reaching (0.09 and 0.19) µA cm^−2^, respectively, before decreasing toward zero. This initial transient current could be attributed to the release of residual electrons stored from the pre-inoculum [51]. Both replicates stabilized on an unsubstantial current of ∼0.02 µA cm^−2^ by the end of the CA. The second replicate started with a current density of 0.1 µA cm^−2^. Over the next 6 h, this current decreased to ∼0.02 µA cm^−2^ and remained below 0.07 µA cm^−2^. The biological nature of this current cannot be confirmed considering its discontinuous and noisy evolution. Overall, these results strongly suggest that *G. sulfurreducens* cannot utilize propionate as sole ED with electrodes serving as EA. While previous results with insoluble Fe(III) oxides did not show propionate consumption in combination with acetate, dedicated experiments will be required to determine if the same stands for propionate oxidation under electrode-respiring conditions.

### Transcriptomic analysis of the propionate-consuming phenotype

To explore the underlying mechanisms of propionate degradation, RNA-seq was conducted on cultures grown with either acetate or propionate as EDs and fumarate as EA, as well as on a ‘shift’ condition generated by adding propionate to the acetate-grown cultures after acetate was fully depleted (Fig. 5A).

**FIG 5.**
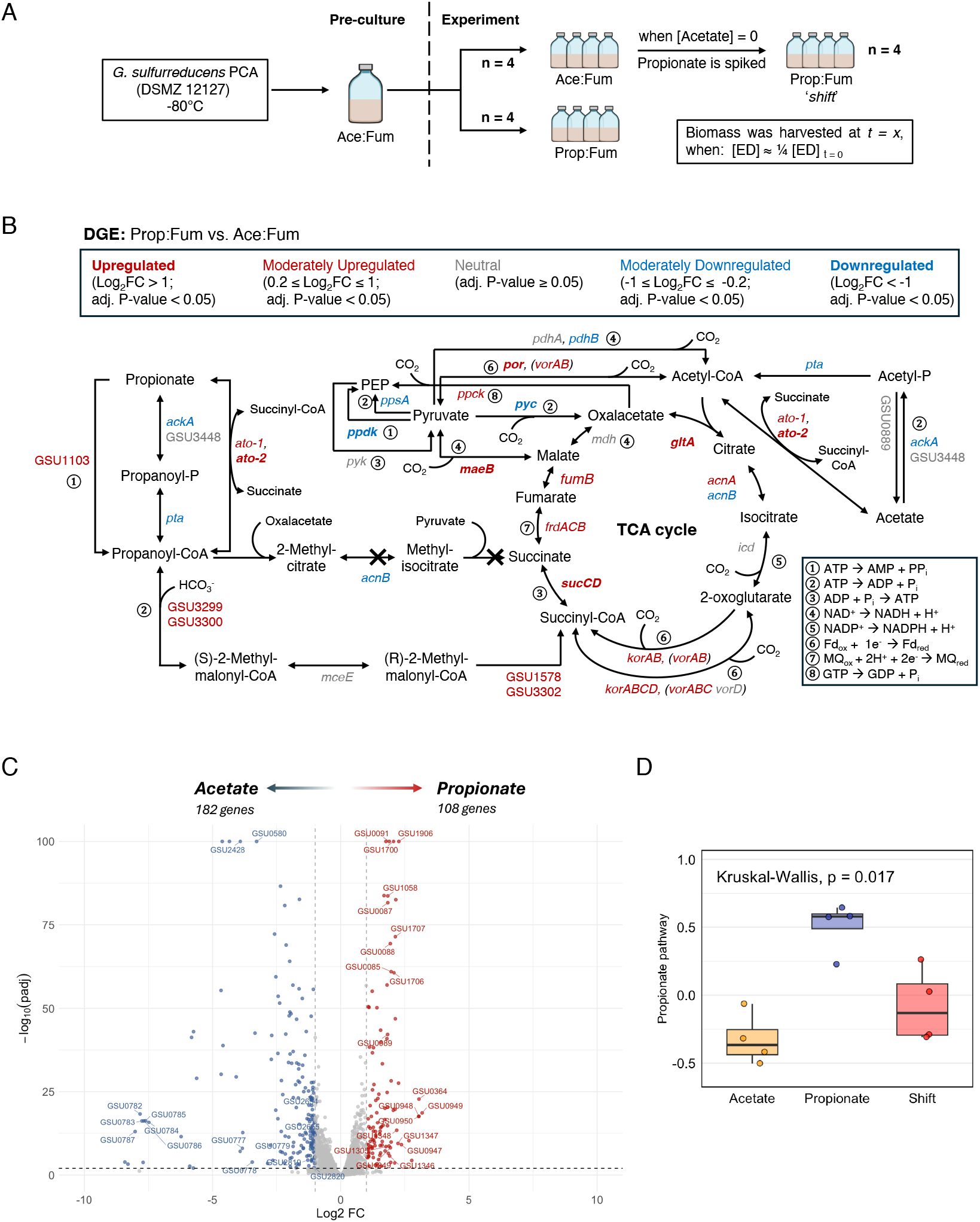
**A)** Diagram of the experimental design for transcriptomic analysis comparing gene expression in *G. sulfurreducens* grown with propionate or acetate as ED. **B)**. DGE analysis between the propionate and acetate grown cultures. The TCA cycle, associated anaplerotic reactions, the acetate degradation pathway, and the putative propionate degradatio_**1**_n_**9**_ pathways are mapped with their respective genes and linked cofactors. Genes are color-coded by adjusted P-value and log_2_FC: grey, no significant differential expression (adj. P-value ≥ 0.05); red, upregulated; blue, downregulated; bold red and bold blue indicate

In *G. metallireducens*, the methylcitrate pathway activates propionate into propanoyl-CoA via a putative succinyl:propionate CoA-transferase (Gmet_1125) and converts it in pyruvate and succinate through 2-methylcitrate intermediates (catalyzed by Gmet_1122–1124) [27]. The MMC pathway forms propanoyl-CoA through an acetate kinase and phosphate acetyltransferase. Then, (S)-2-methylmalonyl-CoA is formed via a biotin-dependent acyl-CoA carboxylase (Gmet_3248-3249), and succinyl-CoA is obtained through an epimerase and mutase [28].

These pathways were considered as potential mechanisms for propionate degradation in *G. sulfurreducens* (Fig. 5B). A local alignment of the amino acid sequences of the proteins involved in propionate degradation from *G. metallireducens* was performed against the *G. sulfurreducens* proteome using the BLASTp tool [52]. The results showed no homologs for key enzymes in the methylcitrate pathway (specifically Gmet_1122 and Gmet_1123). For this reason and due to the lack of evidence in the transcriptomic analysis (*vide infra*), its putative contribution seems unlikely, yet the presence of highly divergent enzymes to serve as functional analogs cannot be dismissed *sensu stricto*. In contrast, protein homologies were found linked to the presence of the MMC pathway. First, the protein encoded by GSU0490 (*ato-1*) shared a 85.9% sequence identity with that of Gmet_1125 (proposed to convert propionate into propanoyl-CoA). Second, the protein products of GSU3299 and GSU3300 exhibited 93.6% and 95.0% identity with Gmet_3248 and Gmet_3249, which are believed to catalyze the conversion of propanoyl-CoA to (S)-methylmalonyl-CoA. Such high identity levels (>80%) strongly suggest conserved enzymatic function between these orthologs [53]. If functional, these genes would support a complete MMC pathway in *G. sulfurreducens* by filling the missing reaction steps noted in the KEGG reference database (accession number ‘*gsu00640*’) [54]; this putative pathway is also not annotated in Gene Ontology (GO) [55] and BioCyc databases [56].

A DGE analysis was performed between the acetate and propionate conditions (Fig. 5C; all results can be found in Extended data 1), identifying 833 genes significantly upregulated in the propionate group (adj. P-value < 0.05), among which, 108 showed substantial upregulation (Log_2_FC *>* 1). Two possible reaction steps for the activation of propionate into propanoyl-CoA showed upregulated genes (Fig. 5B; see the gene list in Table S4). The first involves *ato-2* and the aforementioned *ato-1*. The second option involves GSU1103, which is inferred (Biocyc ID: RS05500) to catalyze its activation with the hydrolysis of ATP to AMP. A third route involving *ackA*, GSU3448, and *pta* may still contribute, yet was downregulated relative to the acetate-only growth condition. Propanoyl-CoA is likely metabolized through the MMC pathway, where a majority of the associated genes were overexpressed. This pathway hydrolyzes ATP and fixes 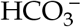, ultimately yielding succinyl-CoA, which enters the TCA cycle [57]. Most TCA cycle genes were upregulated when grown with propionate, except those encoding for *icd* and *mdh*, which were not differentially expressed, and *acnB*, which was instead overexpressed during growth on acetate. *G. sulfurreducens* encodes two aconitase isoenzymes, *acnB* and *acnA*, whose utilization has been linked to exponential growth or stress, respectively [58, 59]. This may suggest that cells grown on propionate are under stress conditions. To obtain a fully functional TCA cycle working with propionate, one molecule of malate is metabolized to oxaloacetate while a second one is converted to acetyl-CoA via the oxidoreductases *maeB* and *por*, both of which showed high transcript levels. Subsequently, acetyl-CoA and oxalacetate are combined to produce citrate via *gltA*, a synthase that was also highly expressed under the propionate condition. As result of a full cycle, energy is generated in the form of ATP, reducing equivalents like NAD(P)H, and reduced ferredoxin (Fd_red_); while at the inner membrane, menaquinone (MQ) is reduced via succinate dehydrogenase (*frd*), contributing to cellular respiration.

In contrast, the acetate group exhibited 788 significantly upregulated genes (adj. P-value < 0.05) with respect to the propionate condition, among which, 182 were strongly upregulated (Log_2_FC < −1; i.e., correspondingly downregulated in the propionate growth condition) (Fig. 5B). Acetate is activated to acetyl-CoA via *ackA* and *pta*. Although *por* is greatly expressed in the propionate growth condition, acetyl-CoA can be transformed to pyruvate via the same pathway using Fd_red_ [60]. Next, pyruvate can be carboxylated into oxalacetate via (*pyc*), which is ATP dependent [61]; as such, the TCA cycle is primed to initiate. It was also observed that the conversion of pyruvate to phosphoenol pyruvate (PEP) was moderately downregulated via *ppsA* in the propionate condition, and strongly downregulated through *ppdk*, suggesting that gluconeogenesis is more active in acetate-grown cultures. Further insights regarding some of the most differentially expressed genes between acetate- and propionate-grown cultures can be found in Supp. Discussion.

To assess the overall activity of the proposed propionate degradation pathway, a GSVA analysis was conducted using 10 candidate genes, namely GSU1103, *ackA, pta, ato-1, ato-2*, GSU3299, GSU3300, *mceE*, GSU1578 and GSU3302. The resulting “propionate signature score” was significantly higher in the propionate condition (adj. P-value < 0.05), supporting the activation of the proposed pathway under these conditions (Fig. 5D). The experimental ‘*shift*’ condition exhibits an intermediate gene expression profile between the acetate and propionate growth conditions indicating a state of transition. Overall, these results support the activity of the MMC pathway for propionate degradation in *G*.*Sulfurreducens*, yet remains to be unequivocally identified.

Considering the activity of the MMC pathway, the thermodynamic estimates presented earlier were recalculated taking into account the energy investment required for propionate metabolism and compared to adjusted values of acetate oxidation (adj. Δ*G*^°′^). Acetate oxidation requires an initial investment of 1 ATP (estimated −56 kJ mol^−1^) for activation to acetyl-CoA. Similarly, propionate degradation requires substrate activation, while additional energy is invested in the carboxylation of propionyl-CoA and the following carbon rearrangement from (R)-methylmalonyl-CoA to succinyl-CoA, resulting in a higher overall energy cost (approximately −116 kJ mol^−1^).

Based on the adj. Δ*G*^°′^ estimates (Table S3), propionate oxidation coupled to Fe(III) citrate provides the lowest energy per electron 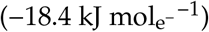 among the conditions tested, whereas fumarate-coupled reactions are energetically more favorable ([−22.6 and −23.9] 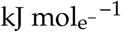 for acetate and propionate, respectively). These results are consistent with the biomass yields obtained experimentally, and places propionate oxidation coupled to Fe(III) citrate reduction close to the thermodynamic boundaries supporting growth in *G. sulfurreducens* [47, 62].

To further investigate the downstream biological effects of propionate degradation in *G. sulfurreducens*, a KEGG-based pathway enrichment analysis was performed on the differentially expressed genes identified under propionate and acetate conditions (Fig. 6).

**FIG 6.**
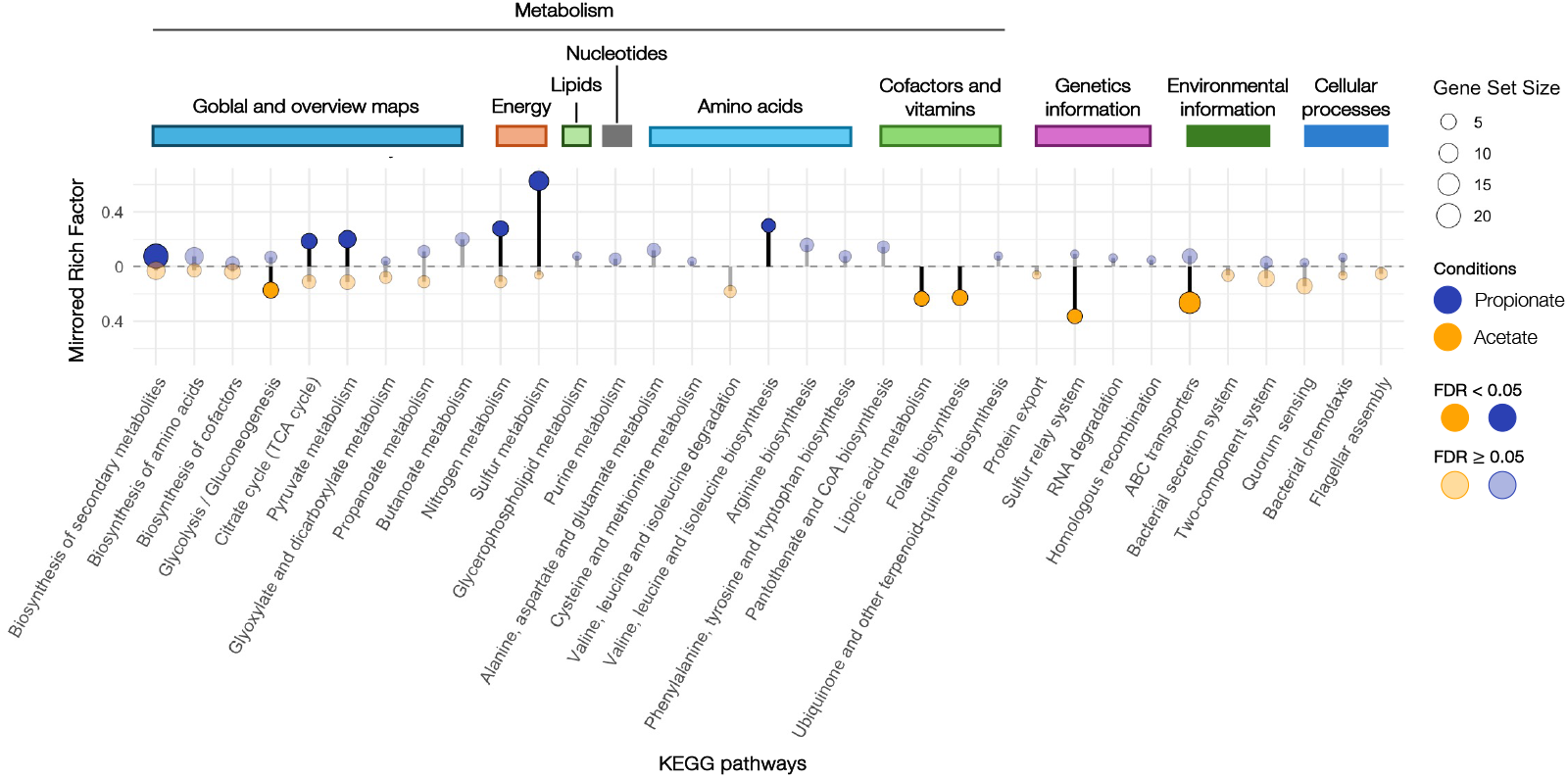
KEGG pathways enriched for propionate and acetate conditions. Circles represent gene sets, with size proportional to the number of genes in each pathway. Upper half correspond to enrichment in propionate (blue), and lower half correspond to enrichment in acetate (orange). Pathways with false discovery rate (FDR) < 0.05 are shown with filled circles.

In the propionate condition, six significantly enriched pathways were identified, including the metabolism of 2-oxocarboxylic acids, the TCA cycle, nitrogen metabolism, sulfur metabolism, and the biosynthesis of branched-chain amino acids (BCAAs). The category of 2-oxocarboxylic acids encompasses key intermediates like pyruvate, oxalacetate, 2-oxoglutarate, and branched-chain keto acids. The concomitant enrichment of such pathways with the TCA cycle is consistent with previous observations and likely reflects an increased flux through the TCA cycle to completely oxidize the three-carbon backbone of propionate to CO_2_. Increased sulfur metabolism under propionate was linked to the biosynthesis of L-cysteine (see Fig. S9A), which acts as the main organic source of sulfur for the assembly of iron–sulfur (Fe–S) clusters [63]. These are essential for redox-active enzymes central to propionate oxidation, including ferredoxin-dependent oxidoreductases, NADH dehydrogenase (complex I), and succinate dehydrogenase. Propionate consumption was also linked to nitrogen metabolism likely due to an increased demand of precursors to sustain biosynthetic pathways. In fact, the synthesis of L-glutamate which serves a central role as amino group donor in the synthesis of amino acids and nucleotides was upregulated (see Fig. S9B) [64]. Consistent with this, the expression of genes associated with biosynthesis of BCAAs (L-valine, L-leucine and L-isoleucine) was significantly enriched (see Fig. S9C). In the context of propionate degradation, which is a more reduced carbon source than acetate, BCAAs can act as temporary carbon sinks, contributing to redox-balancing. Synthesized BCAAs can be incorporated into proteins, be used in membrane lipid synthesis, oxidized to supply reducing power, or be actively exported if accumulated in excess [65]. Notably, propanoyl-CoA can be converted via *korB* (GSU1469) to 2-oxobutanoate (Biocyc ID: RS07310), an intermediate in the synthesis of L-isoleucine. This provides an alternative route for redirecting propionate into anabolism.

The higher energy yield associated to acetate metabolism increased anabolism in acetate-grown cultures. This was reflected by the enrichment of glycolysis/gluconeogenesis and the biosynthesis of lipoic acid and folate (Fig. 6). Lipoic acid is essential in enzymes such as pyruvate and *α*-ketoglutarate dehydrogenases, which are part of the central carbon metabolism and the catabolism of aminoacids [66, 67]. The derivatives of folate supply C1 molecules for the synthesis of nucleotides [68]. Finally, the sulfur relay system, which is involved in the biosynthesis of cofactors and modification of tRNAs [69], as well as ABC transporters, which facilitate nutrient uptake and metabolite export, were enriched [70]. These features highlight the enhanced metabolic activity observed during acetate growth with respect to propionate conditions.

This is the first time, to the best of our knowledge, that *G. sulfurreducens* is shown to oxidize propionate and use it as a carbon source. Propionate was utilized with fumarate as EA, but required the presence of acetate when Fe(III) citrate was the EA. No propionate consumption was observed with Fe(III) oxides or electrodes serving as EAs. The transcriptomic analyses performed strongly support the activity of the MMC pathway in propionate degradation. Other studies have reported the use of propionate in co-cultures of *G. sulfurreducens* with *S. fumaroxidans* [71, 23] or *G. metallireducens* [28] with Fe(III) citrate as EA, yet this capability was never attributed to *G. sulfurreducens*.

Collectively, these results highlight a possible underappreciated prevalence of *Geobacter* species with propionate-degrading capabilities. The utilization of cultures metabolically adapted to oxidizing propionate with fumarate as EA, to test whether these could reduce insoluble EAs is an interesting direction for future studies. Considering the potential relevance of EET-based strategies (DET or DIET) in biotechnological applications to limit propionate accumulation, further investigation of the kinetics involving propionate oxidation by *G. sulfurreducens* is key. Furthermore, pathway-specific gene targeted deletion approaches constitute an immediate next step to unravel the molecular mechanisms supporting propionate metabolism in this species.

## ACKNOWLEDGMENTS

We thank Quinten Mariën, Alberte Regueira, Tom Van de Wiele and Elie Desmond-Le Quéméner for insightful discussions. We acknowledge Jana de Bodt for providing lab support.

## DATA AVAILABILITY STATEMENT

All transcriptomic analysis code scripts and raw sequencing files of bulk RNA-seq have been made available at the GitHub repository: “davidhernandezvillamor-cmyk”. These have also been made available in NCBI SRA (BioProject ID: PRJNA1380372).

## FUNDING

This work was inititated by J.R.B.A in the context of the MELiSSA program (No. 4000137386/22/NL/KML), which is funded by the European Space Agency (ESA). D.H.V is supported by a Bijzonder Onderzoeksfonds grant (BOF21 /DOC /236) from Ghent University. A.P is supported by a fundamental research project from Fonds Wetenschappelijk Onderzoek (G038520N). A.J was supported by the Industrieel Onderzoeksfonds council ‘PACTIOSTAT’ grant (UGent no. F2020/IOF-StarTT/127).

## CONFLICTS OF INTEREST

The authors declare no conflict of interest.

## SUPPLEMENTAL MATERIAL

“20251006_Supp_Information.pdf” “Extended_Data1.xlsx”

## Notes

### Competing Interest Statement

The authors have declared no competing interest.

