## Supporting Information for "Propionate oxidation by *Geobacter sulfurreducens* is electron acceptor dependent"

**<u>Table of Contents:</u>**

Table S1. Summary of replicates per experiment and condition tested.

Table S2. Standard redox potentials and electron stoichiometry for acetate and propionate equations.

Table S3. Standard and adjusted Gibbs free energy changes for propionate and acetate oxidation.

Table S4. Propionate vs. acetate DGE linked to central carbon metabolism in *G. sulfurreducens*.

Figure S1: Precipitation of iron phosphates in modified M9 medium.

Figure S2: Electrochemical set-up to test propionate utilization by *G. sulfurreducens* at anodic plateau potentials.

Figure S3: Growth assay cultures with fumarate as EA and acetate or propionate as the ED.

Figure S4: Cumulative fraction of electrons assumed to go to biomass in propionate and propionate plus acetate cultures using fumarate as EA and evolution of the contribution from propionate.

Figure S5: Growth assay cultures with Fe(III) citrate as EA and either propionate, or propionate and acetate as EDs.

Figure S6: *G. suflurreducens* cell density in iron citrate cultures with acetate plus propionate

Figure S7: Cyclic voltammograms to assess ammonium iron citrate formal potential.

Figure S8: Growth assay cultures with insoluble iron oxides as EA and either propionate, or propionate and acetate as the EDs.

Figure S9: Pathways enriched during propionate consumption with fumarate as the EA.

Supplementary Materials and methods

Supplementary Discussion

Supplementary References

**Table S1.** Replicate counts for each growth assay, split by condition.

| Growth assay | | Replicates | |
|---|---|---|---|
| **Assay** | **Condition** | **Abiotic ctrl.** | **Experiment** |
| Fumarate as EA | Propionate | 2 | 2 |
| | Acetate + Propionate | 2 | 3 |
| soluble Fe(III) citrate as EA | Propionate | 2 | 3 |
| | Acetate + Propionate | 2 | 3 |
| insoluble Fe(III) oxides as EA | Propionate | — | 3 |
| | Acetate + Propionate | — | 3 |
| Glassy carbon electrodes | Acetate | — | 1 |
| | Propionate | — | 3 |
| Fumarate as EA (RNA-seq) | Acetate | — | 4 |
| | Propionate | — | 4 |

**Table S2.** Standard redox potentials and electron stoichiometry for propionate and acetate oxidation with different EAs.

| Reaction | $E^{\circ\prime}_{\text{donor}}$ (V) | $E^{\circ\prime}_{\text{acceptor}}$ (V) | $\Delta E^{\circ\prime}$ (V) | $n$ (e$^-$) |
|---|---|---|---|---|
| Propionate + fumarate | −0.29 | 0.03 | 0.32 | 14 |
| Propionate + Fe(III) citrate | −0.29 | −0.014 | 0.276 | 14 |
| Acetate + fumarate | −0.29 | 0.03 | 0.32 | 8 |
| Acetate + Fe(III) citrate | −0.29 | −0.014 | 0.276 | 8 |

**Table S3.** Standard and adjusted Gibbs free energy changes for propionate and acetate oxidation.

| Reaction | $\Delta G'_{ini}$ (kJ mol$^{-1}$) | $\Delta G^{\circ\prime}$ (kJ mol$^{-1}$) | $\Delta G^{\circ\prime}$ (kJ mol$^{-1}$ e$^{-1}$) | Adj. $\Delta G^{\circ\prime}$ (kJ mol$^{-1}$) | Adj. $\Delta G^{\circ\prime}$ (kJ mol$^{-1}$ e$^{-1}$) |
|---|---|---|---|---|---|
| Propionate + fumarate | −477.7 | −432.32 | −30.88 | −316.32 | −22.59 |
| Propionate + Fe(III) citrate | −376.8 | −372.88 | −26.63 | −256.87 | −18.35 |
| Acetate + fumarate | - | −247.04 | −30.88 | −191.04 | −23.88 |
| Acetate + Fe(III) citrate | - | −213.07 | −26.63 | −157.07 | −19.63 |

**Table S4.** Comparison of central carbon metabolism gene expression levels in *G. sulfurreducens* grown on propionate vs. acetate, categorized by $Log_2FC$ values and statistical significance.

| Gene ID | Gene (abbr.) | Full Name | $Log_2FC$ |
|---|---|---|---|
| **adj. P-value $Log_2FC < 0.05$, Downregulated ($Log_2FC < 0$)** | | | |
| GSU2707 | *ackA* | acetate kinase | -0.86 |
| GSU2706 | *pta* | phosphoacetyl transferase | -0.48 |
| GSU2428 | *pyc* | pyruvate carboxylase | -3.92 |
| GSU0580 | *ppdk* | pyruvate phosphate dikinase | -3.28 |
| GSU0803 | *ppsA* | phosphoenolpyruvate synthase | -0.88 |
| GSU2436 | *pdhB* | pyruvate dehydrogenase beta subunit | -0.41 |
| GSU1660 | *acnB* | aconitate hydratase 2 | -0.64 |
| **adj. P-value $Log_2FC < 0.05$, Upregulated ($Log_2FC > 0$)** | | | |
| GSU0490 | *ato-1* | acetyl-CoA hydrolase 1 | 0.88 |
| GSU0174 | *ato-2* | acetyl-CoA hydrolase 2 | 1.39 |
| GSU846 | *acnA* | aconitate hydratase 1 | 0.49 |
| GSU1058 | *sucC* | succinyl-CoA synthetase beta subunit | 1.83 |
| GSU1059 | *sucD* | succinyl-CoA synthetase alpha subunit | 1.69 |
| GSU1176 | *frdC* | succinate dehydrogenase/fumarate reductase (cytochrome b558 subunit) | 0.54 |
| GSU1177 | *frdA* | succinate dehydrogenase/fumarate reductase (flavoprotein subunit) | 0.68 |
| GSU1176 | *frdB* | succinate dehydrogenase/fumarate reductase (iron-sulfur protein) | 0.66 |
| GSU0994 | *fumB* | fumarate hydratase | 0.80 |
| GSU1106 | *gltA* | citrate synthase | 1.24 |
| GSU1468 | *korA* | 2-oxoglutarate ferredoxin oxidoreductase (alpha subunit) | 0.73 |
| GSU1469 | *korB* | 2-oxoglutarate ferredoxin oxidoreductase (ThDP-binding subunit) | 0.73 |
| GSU1470 | *korC* | 2-oxoglutarate ferredoxin oxidoreductase ((gamma subunit)) | 0.73 |
| GSU1467 | *korD* | 2-oxoglutarate ferredoxin oxidoreductase (ferredoxin subunit) | 0.43 |
| GSU1861 | *vorA* | 2-oxoglutarate ferredoxin oxidoreductase (alpha subunit) | 0.40 |
| GSU1860 | *vorB* | 2-oxoglutarate ferredoxin oxidoreductase (ThDP-binding subunit) | 0.34 |
| GSU1859 | *vorC* | 2-oxoglutarate ferredoxin oxidoreductase (gamma subunit) | 0.41 |
| GSU1700 | *maeB* | malate oxidoreductase | 2.06 |
| GSU3385 | *ppck* | phosphoenolpyruvate carboxykinase | 0.84 |
| GSU0097 | *por* | pyruvate:ferredoxin/flavodoxin oxidoreductase | 1.23 |
| GSU1103 | - | fatty-acyl-CoA synthase | 0.39 |
| GSU3299 | - | biotin-dependent acyl-CoA carboxylase (carboxyltransferase subunit) | 0.35 |
| GSU3300 | - | biotin-dependent acyl-CoA carboxylase (biotin carboxylase subunit) | 0.41 |
| GSU1578 | - | (R)-methylmalonyl-CoA mutase (adenosylcobamide-binding subunit) | 0.44 |
| GSU3302 | - | (R)-methylmalonyl-CoA mutase (isobutyryl-CoA mutase-like catalytic subunit) | 0.38 |
| **adj. P-value $\geq 0.05$ (Not Significant)** | | | |
| GSU3303 | *mceE* | methylmalonyl-CoA epimerase | 0.29 |
| GSU3331 | *pyk* | pyruvate kinase | 0.16 |
| GSU3448 | - | acetate kinase-related protein | – |
| GSU1862 | *vorD* | 2-oxoglutarate ferredoxin oxidoreductase (ferredoxin subunit) | – |
| GSU1465 | *icd* | isocitrate dehydrogenase | 0.26 |
| GSU2443 | *pdhA* | pyruvate dehydrogenase alpha subunit | -0.12 |
| GSU0889 | - | acylphosphatase | -0.25 |
| GSU1466 | *mdh* | malate dehydrogenase | 0.17 |

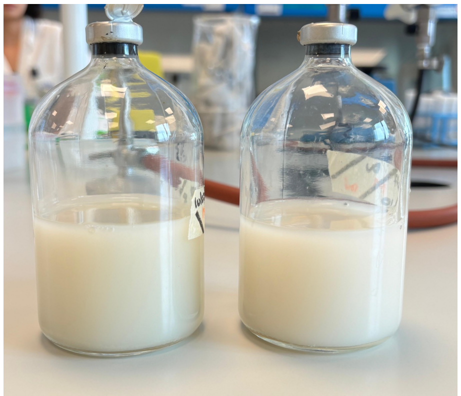

Fe(II)-phosphates formation

**Fig. S1.** Cultures of *G. sulfurreducens* grown in modified M9 medium showing the formation of iron phosphates.

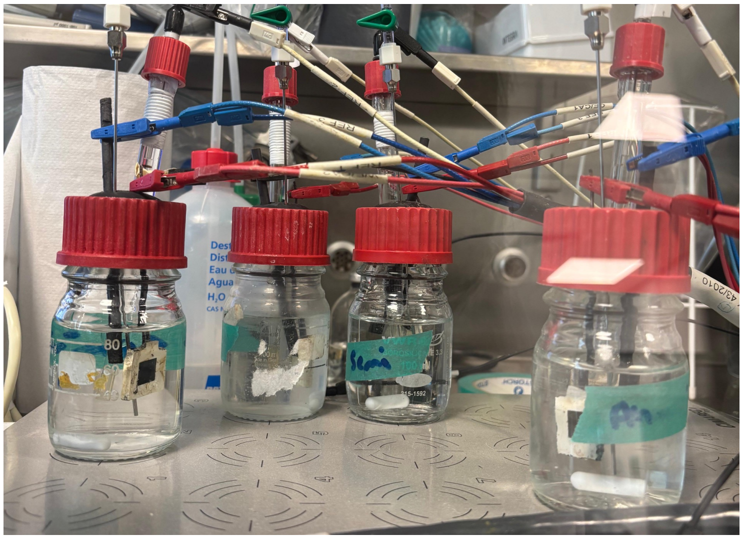

**Fig. S2.** Electrochemical setup enclosed in an anerobic chamber. It comprises four 100 mL bottles placed over a common stirring plate, each equipped with a glassy carbon working electrode (poised at +0.1 V vs. SHE), a graphite rod as counter electrode and a Ag/AgCl 3M KCl reference electrode.

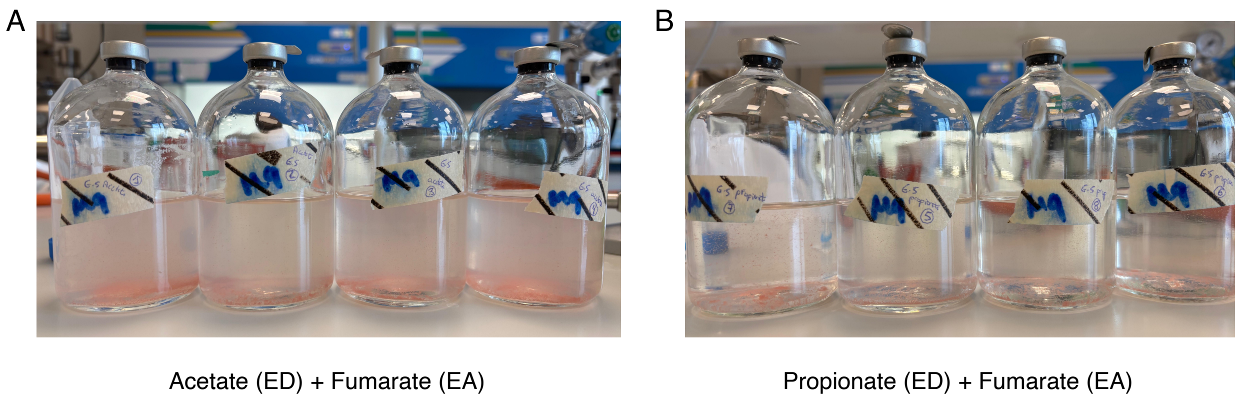


**Fig. S3.** Stationary-phase cultures of *G. sulfurreducens* grown with acetate **(A)** or propionate **(B)** as the only ED, and fumarate as the EA.

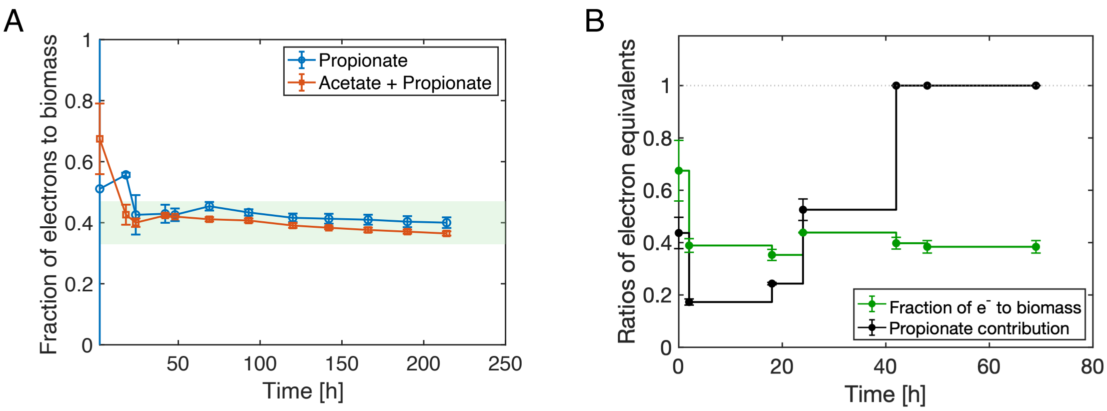


**Fig. S4. (A)** Cumulative fraction of electrons assumed to go to biomass and microbial reducing equivalent i.e. electrons not ending up in succinate through respiration; for propionate as ED (blue) and propionate and acetate (orange). The green band shows the range of values reported in the literature for *G. sulfurreducens* grown with acetate and fumarate, averaging at 40 ± 7% (six conditions from three studies [1-3]). **(B)** Sequential evolution of this electron yield for the growth with acetate and propionate, exhibited for each period between two successive measurements, independently (green); the black curve shows the relative contribution of propionate as ED for all sequential periods.

A

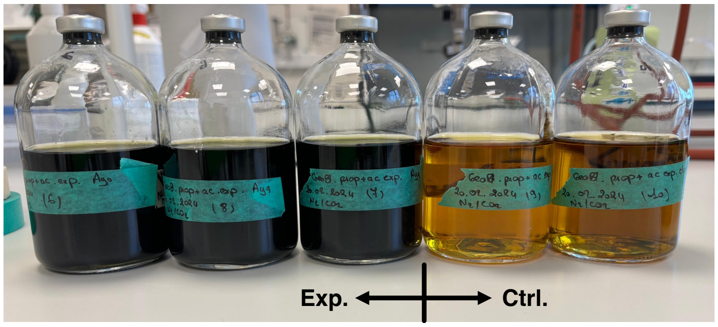


Propionate + Acetate (EDs) + sol. Fe(III) citrate (EA)

B

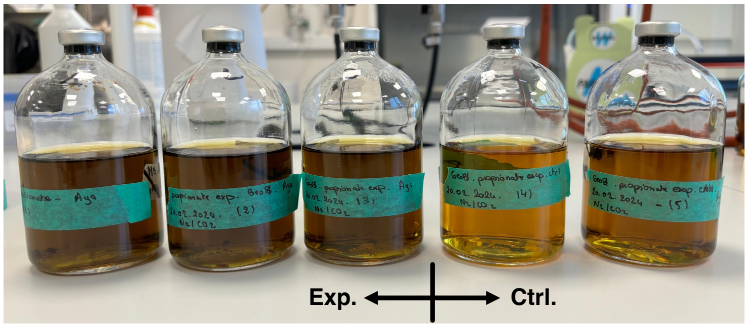


Propionate (ED) + sol. Fe(III) citrate (EA)

**Fig. S5.** Stationary-phase cultures of *G. sulfurreducens* grown with acetate plus propionate **(A)** or propionate alone **(B)** as the EDs, and soluble Fe(III) citrate as the EA. Abiotic controls are shown on the right-hand side of each image.

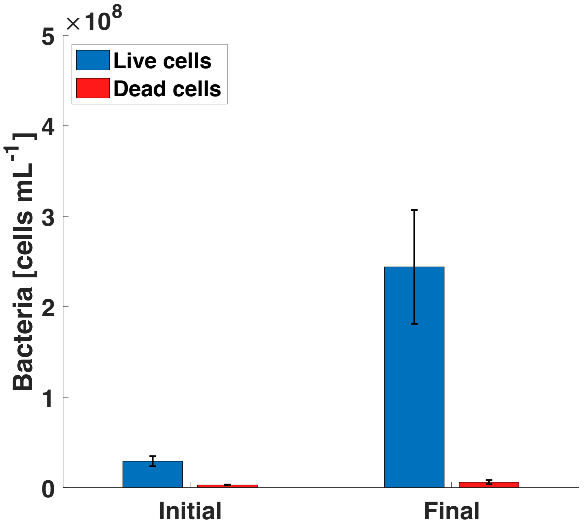


**Fig. S6.** *G. sulfurreducens* cell density in iron citrate cultures with acetate plus propionate at the beginning and end of the experiment.

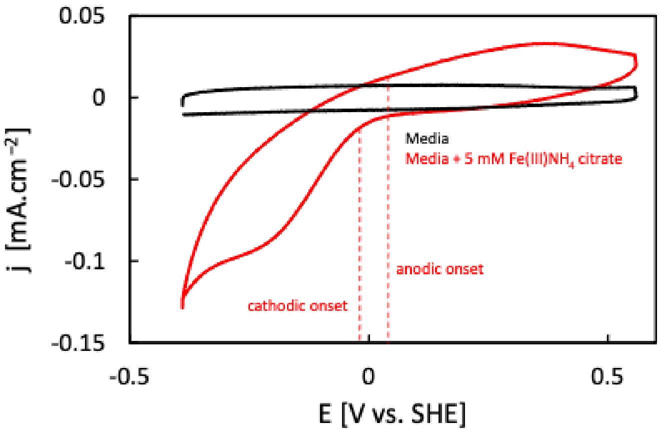


**Fig. S7.** Cyclic voltammograms recorded at 20 mV $s^{-1}$ in (black) medium and (red) medium with 5 mM of ammonium iron citrate in anaerobic conditions. Working electrode: polished glassy carbon; counter: platinum wire; reference: Ag/AgCl (3 M KCl). The CVs are not typical of reversible electron transfer, and it is therefore impossible to obtain a precise formal potential. Yet, it is reasonable to assume that the formal potential lies between the reduction and oxidation onsets.

The $E^{\circ\prime}$ value from Fe(III) citrate has been reported to range from (~0 to 0.37) V vs. SHE at circumneutral pH [4-6]. To account for this variability, cyclic voltammetry was performed in the media utilized during Fe(III) assays, both in the absence and presence of ammonium iron citrate, which displayed quasi-irreversible electron transfer behavior. Still, the reduction and oxidation onsets observed are in line with a $E^{\circ\prime}$ value close to 0 V vs. SHE, thus remaining comparable to that of fumarate ($E^{\circ\prime}$ = +0.03 V vs. SHE) [7].

A B

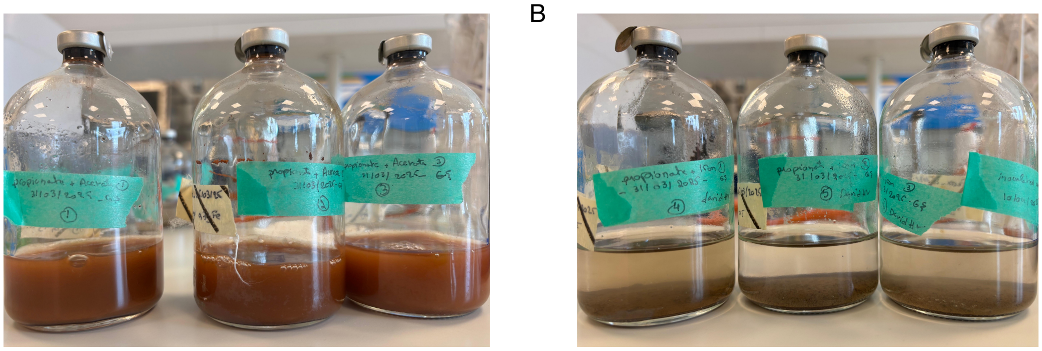

Propionate + Acetate (EDs) + insol. Fe(III) oxides (EA) Propionate (ED) + insol. Fe(III) oxides (EA)

**Fig. S8.** Stationary-phase cultures of *G. sulfurreducens* grown with acetate plus propionate **(A)** or propionate alone **(B)** as the EDs, and insoluble Fe(III) oxides as the EA.

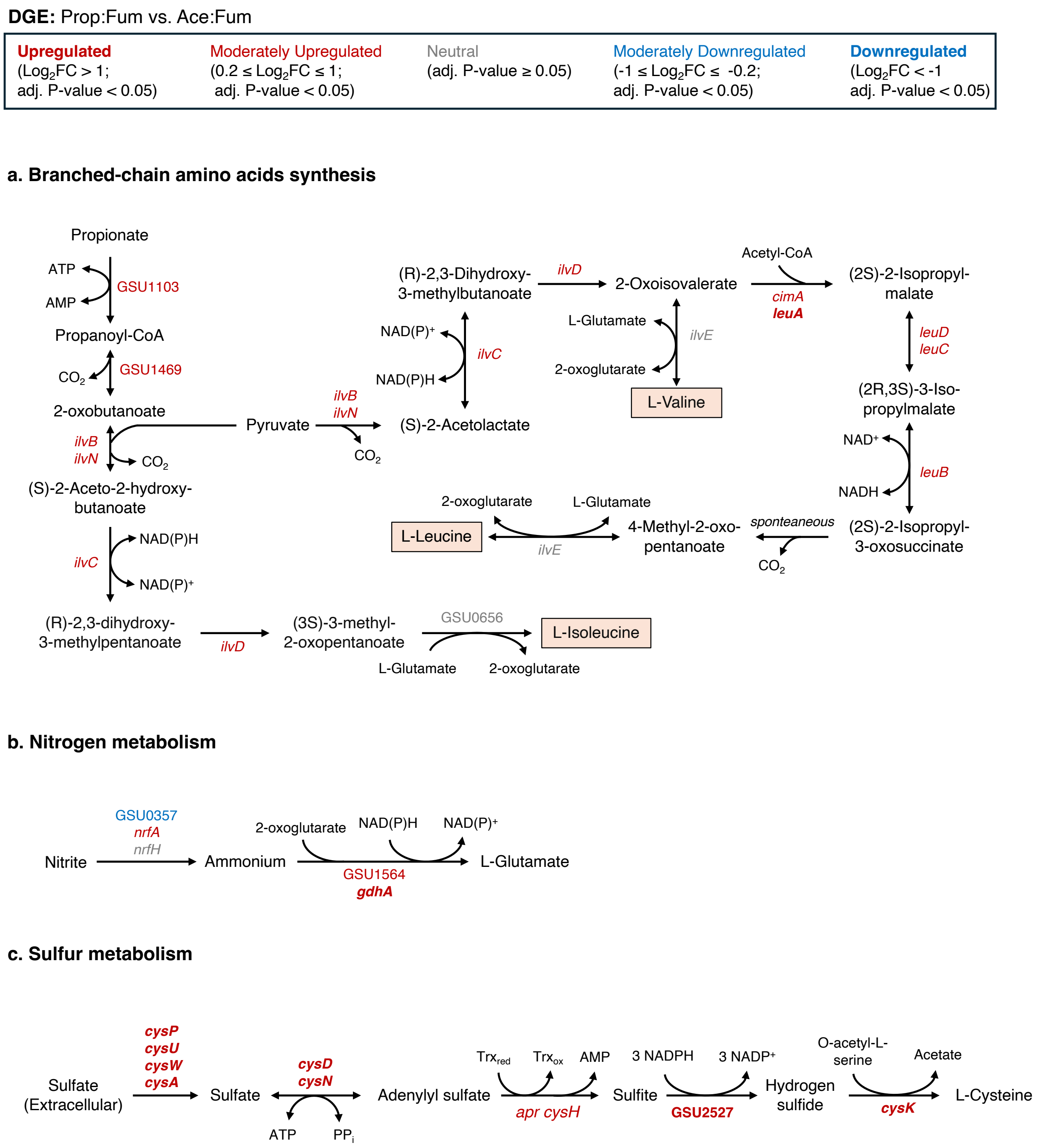


**Fig. S9.** Metabolic pathways based on DGE analysis between propionate and acetate grown cultures of *G. sulfurreducens*. (A) BCAAs biosynthesis pathway, (B) nitrogen metabolism toward L-glutamate biosynthesis and (C) sulfur metabolism toward L-cysteine biosynthesis. Associated genes are color-coded by adjusted P-value and $\log_2$FC: grey, no significant differential expression (adj. P-value ≥ 0.05); red, upregulated; blue, downregulated; bold red and bold blue indicate strong upregulation ($Log_2FC > 1$) and strong downregulation ($Log_2FC < -1$), respectively.

### Supplementary Materials and methods

#### Ion chromatography

*G. sulfurreducens* cells were removed from organic acid samples by filter sterilization, then analyzed using a Compact IC Flex (Metrohm) with inline bicarbonate removal (MCS). An 858 Professional Sample Processor with extended MiPuT equipment (Metrohm) was used for sample injection. A Metrosep organic acids (250/7.8) and Metrosep organic acids guard column (Guard/4.6) were used for compound separation, and an 850 IC conductivity detector was used to detect eluted components.

#### Flow cytometry

Samples were diluted with sterile phosphate-buffered saline (PBS) and incubated in darkness at 37℃ for 20 min with a viability staining mix of SYBR® Green I (100x concentrate, Invitrogen) and propidium iodide (50x, 20 mM concentrate, Invitrogen) in 0.22 µm-filtered dimethyl sulfoxide (Sigma Aldrich) for live-dead analysis. Subsequently, the samples were analyzed immediately using an Attune NxT BVXX (Thermo Fisher Scientific, Belgium) equipped with eight fluorescence detectors: 530/30 nm, 574/26 nm, 695/40 nm, 780/60 nm (Blue Channel), 440/50 nm, 512/25 nm, 603/48 nm, 710/50 nm (Violet Channel), and two scatter detectors, a 50 mW 488 nm blue laser and a 50 mW 405 nm red laser. Heat killed samples (80°C for 1 h), 0.22 µm-filtered samples and sterile PBS were included to assist in the gating of the damaged population of cells and identifying background noise.

#### Electrochemical data treatment

EC-lab® software data files were transformed to '.mpt' format utilizing built-in functionalities, then converted to '.txt' and further analyzed using MATLAB® R2021a, from which CA graphs were derived.

#### DNA extraction protocol and sequencing

*G. sulfurreducens* cultures were confirmed axenic via Sanger sequencing. Samples were mixed with a lysis buffer consisting of 100 mM Tris pH8, 100 mM EDTA pH 8.0, 100 mM NaCl, 1% polyvinylpyrrolidone (PVP40) and 2% sodium dodecyl sulphate (SDS). Glass beads of 0.1 mm diameter were added to the samples. Cells were disrupted using BioLyzer (MoBio) at 2000 rpm for 300 s. The samples were centrifuged for 5 min at 19,000 g, then the supernatant was added to a new tube containing 500 µL of phenol:chloroform:isoamilic alcohol 25:24:1 at pH 7. Samples were mixed, then centrifuged and the resulting upper aqueous phase was drawn to a new tube containing 700 µL of chloroform. Samples were mixed, then centrifuged, and 450 µL of the upper phase was added to a new tube containing 500 µL of cold isopropanol and 45 µL of 3 M sodium acetate. The samples were mixed and stored at -20 °C for one hour after which they were centrifuged at 4 °C for 30 minutes. The supernatant was removed and the DNA pellet dried prior to dissolving in 1X TE buffer. Amplification of highly conserved bacterial 16S rRNA gene was performed by PCR using primers: 27F, 5'-GAGTTTGATCMTGGCTCAG-3' in the forward direction and

1492R, 5’-GGYTACCTTGTTACGACTT-3’ for the reverse. Correct length PCR fragments were purified using GeneJET Gel Extraction Kit (ThermoFisher Scientific); >10 ng DNA $\mu L^{-1}$ were ensured and sent for Sanger sequencing (LGC genomics GmbH).

### Supplementary Discussion

Several of the most highly expressed genes in the propionate condition (Fig. 5C) included the *hdr* locus (GSU0085–GSU0091), annotated by UniProt as a heterodisulfide oxidoreductase system. These systems have not only been associated with the reduction of sulfate, but also with the oxidation of organic compounds [8] and participation as electron bifurcation mechanisms to regenerate reduced ferredoxin, conserve energy, and perform redox balance in anaerobic metabolisms [9, 10]. The *nuo-1* locus (GSU0338–GSU0351), which encodes the NADH:quinone oxidoreductase complex I, was also upregulated, along with *panB* (GSU1705) and *panC* (GSU1706), which are involved in the biosynthesis of pantothenate linked to Coenzyme A production. In contrast, the *nuo-2* locus (GSU3429–GSU3436, GSU3439, GSU3441–GSU3445), a paralog of *nuo-1*, was downregulated. Notably, *ppcB* (GSU0364), which encodes for a periplasmic cytochrome c, was the second most upregulated gene, highlighting a possible role in the electron transport chain during propionate metabolism. Additionally, GSU0947–GSU0950 were among the most strongly upregulated genes. The first two are predicted ABC transporters, while the second pair encodes RND (resistance-nodulation-cell) system-like efflux pumps. Both systems may also contribute to propionate transport and detoxification of metabolites formed during its metabolism [11].

Membrane-bound [NiFe] hydrogenase genes (*hyb*, GSU0782- GSU0787) ranked within the 10 most upregulated genes during acetate consumption (Fig. 5C). These are known to produce $H_2$ coupled to the oxidation of cytochromes, $Fd_{red}$, and NAD(P)H [12], thus suggest that hydrogen cycling contributes more significantly to redox balancing or energy conservation during acetate metabolism. Consistent with this, previous studies have reported traces of $H_2$ (~ 50 Pa) in the gas phase of *G. sulfurreducens* cultures growing on acetate as ED [13, 14]. In contrast, the expression of these genes was downregulated (> 36-fold) when the species was co-cultured with *Syntrophobacter fumaroxidans* [15], and when cultured axenically under different EDs, the expression of *hyb* associated genes was highest with formate, followed by $H_2$ and acetate [16]. This would theoretically place the propionate condition tested here at the lowest level of *hyb* gene expression, which is consistent with a minimal involvement of hydrogen under these growth conditions.
